# A statistical approach to identify regulatory DNA variations

**DOI:** 10.1101/2023.01.31.526404

**Authors:** Nina Baumgarten, Laura Rumpf, Thorsten Kessler, Marcel H. Schulz

## Abstract

Non-coding variations located within regulatory elements may alter gene expression by modifying Transcription Factor (TF) binding sites and thereby lead to functional consequences like various traits or diseases. To understand these molecular mechanisms, different TF models are being used to assess the effect of DNA sequence variations, such as Single Nucleotide Polymorphisms (SNPs). However, few statistical approaches exist to compute statistical significance of results but they often are slow for large sets of SNPs, such as data obtained from a genome-wide association study (GWAS) or allele-specific analysis of chromatin data.

**Results:** We investigate the distribution of maximal differential TF binding scores for general computational models that assess TF binding. We find that a modified Laplace distribution can adequately approximate the empirical distributions. A benchmark on *in vitro* and *in vivo* data sets showed that our new approach improves on an existing method in terms of performance and speed. In applications on large sets of eQTL and GWAS SNPs we could illustrate the usefulness of the novel statistic to highlight cell type specific regulators and TF target genes.

**Conclusions:** Our approach allows the evaluation of DNA changes that induce differential TF binding in a fast and accurate manner, permitting computations on large mutation data sets. An implementation of the novel approach is freely available at https://github.com/SchulzLab/SNEEP.

## 1 Introduction

Non-coding Single Nucleotide Polymorphisms (SNPs) localized in regulatory elements such as enhancers or promoters can lead to changes in gene expression by modifying Transcription Factor Binding Sites (TFBS). Several studies reported the resulting functional consequences (reviewed for instance in [51]). Therefore, methods pinpointing to such regulatory SNPs (rSNPs) are a topic of current interest.

Several methods exist that successfully highlight rSNPs based on epigenetic information like open chromatin data, TF and histone ChIP-seq without taking into account which TF might be affected [30,25,2,12]. In contrast, methods that evaluate the effect of a SNP on a TFBS rely on the ability to describe the binding behaviour of a Transcription Factor (TF) to assess the difference induced by a non-coding SNP. TFs are able to bind the DNA by recognizing short patterns. These patterns can be described *in vitro* using high throughput methods like protein-binding microarrays (PBM) [7] or SELEX [22], or in vivo using ChIP-based techniques [29,21]. The identified TF binding preferences are summarized in a TF model, most prominent are Position Weight Matrices (PWMs) [45]. However, there are other more complex TF models utilizing a bayesian network or other types of Markov models (reviewed in [8]). Other proposed models are for instance the binding energy model (BEM) [53] or the transcription factor flexible model (TFFM) [35], the latter being available within the JASPAR database [11]. Further there is the SLIM model [24], which additionally provides a graphical visualisation similar to the sequence logos of PWMs, and methods based on deep convolutional neural networks like DeepBind [1] or BPNet [4].

Computational approaches have been developed to evaluate the effect of a SNP on the binding sites of a TF. Among them is for instance *GERV* [50], a *k*-mer based approach that learns *de novo* TF binding based on open chromatin and TF ChIP-seq data. To evaluate the impact of a SNP they compute the difference of the predicted read counts for the two allelic variants of a SNP. A more recently published method is *FABIAN-variants* [44]. They determine a differential TF binding score not only based on PWMs but also on TFFMs and allow to take TFBS from epigenetic data into account. In comparison, QBIC-Pred [34,52] utilizes *in vitro* universal PBM data to determine the TF binding behaviour with a *k*-mer based model using ordinary least squares (OLS). They score the effect of a SNP based on the parameters of the OSL z-score, and additionally provide a *p*-value for their score. Further, there are methods like *rTRAP* [33], *is-rSNP* [32] or *atSNP* [54] that rely solely on TF models (usually PWMs) and the DNA sequence itself. To evaluate the effect of a SNP on TF binding sites different statistical approaches are introduced by these methods. *rTRAP* ranks TFs based on the difference of the TF binding scores for the wildtype and the alternative allele of a SNP, whereas *is-rSNP* and *atSNP* provide a *p*-value for their differential TF binding score.

In general, we noticed that only a few methods provide a statistical significance for their introduced differential TF binding scores. However, this is necessary to decide if a score is significantly different from the commonly assumed null hypothesis that the SNP does not affect the TF binding site. Further if the TF binding score is represented as *p*-value they are directly comparable between the TFs, which is often not possible for the scores itself. When the methods provide a *p*-value, their statistic is dependent on their underlying TF model. For instance, QBIC-Pred derives their test statistic based on OLS estimation of *k*-mers, whereas *atSNP* assumes the scores follow a multinomial distribution to model PWMs. *is-rSNP* is with the best of our knowledge the only method which allows to compute a *p*-value independent from the TF model. However, they determine the exact *p*-value distribution for differential TF binding scores by assessing all possible single base changes resulting in a quadratic algorithm prohibitive for large data sets.

In this work, we introduce a fast and accurate approach to provide statistical significance for the differential TF binding score for general TF models. To do so, we examine the distribution of the maximal differential TF binding scores and find that it can be well approximated by a modified Laplace distribution. By using the modified Laplace distribution, we can derive a *p*-value for the maximal differential TF binding score in constant time. We show on experimentally validated TF-SNP pairs that our approach improves the performance in comparison to the previously established method *atSNP*, while being an order of magnitude faster. As applications we present the identification of cell type specific TFs whose binding sites are perturbed by eQTLs in lymphocytes and fibroblasts. Further, we showcase how to combine our approach with publicly available regulatory elements (REMs), derived from epigenomics data, to pinpoint potential target genes affected by rSNPs in an atherosclerosis GWAS.

## 2 Methods

### 2.1 Definition of the problem

In the following, we explain how to evaluate the effect of a SNP on a TFBS. In the first problem definition we explain how to compute the differential TF binding score for a fixed TF position. The second definition describes a more general case where the differential TF binding is computed for all sequences overlapping the SNP.

#### The differential TF binding problem

Let *M* be a general TF model of length *m, S* = {*s*_1_,…, *s_m_*} a DNA sequence with *s_i_* ∈ Σ over the alphabet Σ = {*A,C,G,T*}, also of length *m* and assume there is a SNP with two allelic variants called wildtype and alternative allele at position *i* with *i* ∈ {1, …,*m*} in *S*. Hence, we consider two variants of the sequence *S, S*^1^ containing the wildtype allele and *S*^2^ the alternative allele. For each of these sequence variants we compute a TF binding score describing the TF binding affinity to the sequence according to the model *M*. From the distribution of all binding affinity values for a TF under the model *M*, we can compute the probability *P*(*a* ≤ *t, M*), the *p*-value of observing a binding affinity *a* smaller than a threshold t. We can use this to determine the corresponding *p*-value of the model *M* for each variant, given as *p*(*S*^1^, *M*) and *p*(*S*^2^, *M*), respectively. The exact computation of the TF binding score and the *p*-value depend on the used TF model (see Methods 2.4). To evaluate the effect of a SNP on a TFBS, we compute a log-ratio similar to Manke *et al.* [33] called *differential TF binding score (D)*:

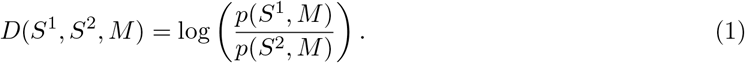

A positive *D* indicates that the exchange from the wildtype allele to the alternative allele increases the binding affinity of the TF and may lead to a gain of a binding site. Whereas a negative D decreases the binding affinity and therefore the binding site might be lost.

#### The maximal differential TF binding problem

Usually it is not known at which position the binding affinity is most affected by the SNP. As a consequence, we evaluate all sequences overlapping with the SNP, hence we define a window of size 2*m* – 1 centered around the SNP. We slide the TF model over the 2*m* – 1 long sequence and compute *D* for each of the sub-sequences of length *m* overlapping the SNP. To identify the optimal *D*, we keep the maximal absolute value.

Therefore, our definition changes: We are given a DNA sequence *S* = {*s*_1_,…, *s*_2*m*-1_} and a SNP with two allelic variants centered in the middle of the sequence at position *m*. We consider two variants of the sequence *S*: *S*^1^ containing the wildtype allele and *S*^2^ including the alternative allele. Further, a *k*-mer which is a sub-sequence of S of length *m*, is defined as *k_i_* = {*s*_1_,…, *s*_*i*+*m*-1_} with *i* ∈ {1,…,*m*}. For each sequence variant, we define the *k*-mers overlapping the SNPs. The *k*-mers of *S*^1^ are denoted as 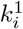 and the *k*-mers for *S*^2^ as 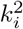 with *i* ∈ {1, …,*m*}. We maximise over the absolute *D* values of all *k*-mers overlapping the SNP and identify the TF position where the binding site is most affected. The *absolute maximal TF binding score D_max_* is defined as:

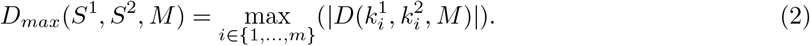

### 2.2 Estimating the distribution of differential TF binding scores

In this section, we investigate if the distribution of *D* follows a known distribution. To do so, we describe the *p*-values of the TF binding scores for the sequences *S*^1^ and *S*^2^ of length *m* as random variables *W* and *Z*, respectively. Since the TF binding score itself is continuous, the *p*-values are uniformly distributed in [0,1] under the null hypothesis [47]. Further, we assume that the random variables W and Z are independent from each other. Hence, the following theorem can be applied:

#### Theorem 1.

*If two independent random variables W, Z, are uniformly distributed in the range* [0,1] *then* 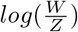 *is Laplace*(0,1) *distributed [27, p. 24]*.

In theory, we conclude that *D* can be represented as random variable 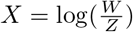 which is *Laplace*(0,1) (*L*(0,1)) distributed. However, our experiments will show that *D* is not following the *L*(0,1) distribution (see Results 3.1). Nevertheless, *D* can be approximated by a *L*(0, b) distribution, with a scale parameter *b* different from 1. The scale parameter is given as 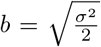, where *σ*^2^ is the variance of the observed differential TF binding scores of a TF model *M* for a set of SNPs.

### 2.3 Derivation of the distribution of the maximal differential TF binding scores

When computing *D_max_,* we can represent the differential TF binding scores of the *k*-mers as *n* independent and identical *L*(0, *b*) distributed random variables *X_i_* with *i* ∈ {1, …,*n*}, where *n* is the overall number of *k*-mers. *D_max_* can be described as the absolute maximum over all random variables *X_i_*, so *Y* = max_*i*∈_{*i*,…,*n*}(|*X_i_*|). To efficiently compute a *p*-value for *D_max_,* we are interested in identifying the cumulative distribution function (CDF) of *Y*.

To do so, we split the following in three parts: First we determine the probability distribution function (PDF) and CDF of the absolute values of *n* Laplace(0,*b*) distributed variables. Second, we derive the CDF of the *n* maximal *L*(0, *b*) distributed random variables, and finally, we combine both parts to get the CDF of *Y*.

#### Computation of the PDF and CDF of the absolute values of *n L*(0, *b*) distributed random variables

The general PDF of the *L*(0, *b*) distribution with the scale *b* > 0 is defined as

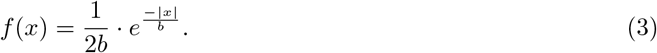

To obtain the density for the absolute value |*x*|, one needs to add up the densities for the positive and negative values:

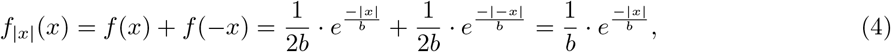

with *x* ∈ ℝ^+^. The corresponding CDF is given by integrating *f*_|*x*|_ (*x*) from 0 to *x*:

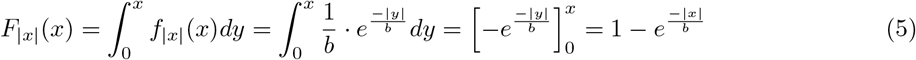

#### Derivation of the CDF of *n* maximal *L*(0, *b*) distributed random variables

The CDF of a random variable *V* is defined as *F*(*x*) = *P*(*V* ≤ *x*). We derive the CDF of the maximal *X_i_* with *i* ∈ {1, …,*n*} using the definition of CDFs and the independence of the *n L*(0, *b*) distributed random variables:

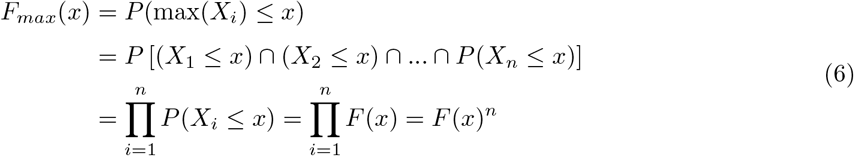

#### Derivation of the CDF for an absolute maximal *L*(0, *b*) distributed random variable

To finally get the CDF of *Y* = max_*i*∈_{1,…,*m*}(|*X_i_*|), we can plug in the CDF for the absolute *L*(0, *b*) distributed random variables (5) in Equation 6 resulting in:

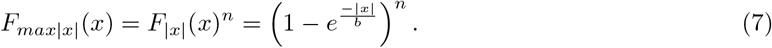

The corresponding PDF is the derivative of equation 7:

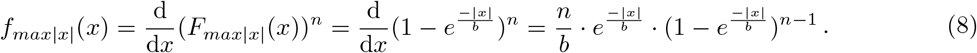

To sum up, we are able to mathematically describe the distribution of *D_max_* as follows:

#### Theorem 2.

*Let D_max_ be defined as Y* = max_*i*∈_{1,…*n*}(|*X_i_*|), *where the X_i_’s are independent and identical L*(0, *b*) *distributed random variables, then Y follows a modified Laplace distribution L_max_*(*n, b*) = *f*_*max*|*x*|_(*x*).

The distribution of *D_max_* is depending on the parameter *n* and the scale parameter *b*. Here, *n* denotes the number of *k*-mers overlapping the SNP times two, since we also consider the reverse complement. To determine the scale parameter *b* for *j* observed maximal differential TF binding scores *x_i_* with *i* ∈ {1,…, *j*} of a TF model *M*, we set up a maximum log-likelihood estimator (MLE) of the PDF of *L_max_*

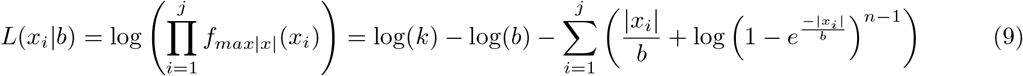

and computed the corresponding derivative with respect to *b*. Since the resulting equation is not analytically solvable we used Newton’s method to numerically approximate *b* (details in Supp. Sec. S1).

Using the CDF of the *L_max_*(*n, b*) distributed maximal differential TF binding scores, we are able to compute a *p*-value for *D_max_* as 1 – *F*_*max*|*x*|_(*x*).

### 2.4 Computation of differential TF binding scores with position weight matrices

Throughout this manuscript, we illustrate our statistical approach using position weight matrices (PWMs) as an example TF model. A PWM *M* describing a TF motif of length *m,* is a 4 × *m* matrix, holding for each base the log-likelihood at position *i* with *i* ∈ {1,…, *m*}. The TF binding score for a sequence *S* given a PWM *M* is computed as 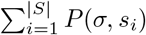, where |*S*| = *m* and *σ* is the base in *S* at position *i*. Using the dynamic programming approach from Beckstette *et al.* [6] we compute for a PWM *M* the exact TF binding score distribution (see Supp. Sec S2). As a result, we can look up the *p*-value for every possible TF binding score obtainable by the PWM *M*. If the *p*-value of a TF binding score is smaller than a given threshold *t,* we assume that the TF, represented by the PWM, is able to bind to the sequence.

### 2.5 Collecting allele specific binding events

We collected 1.760 Allele Specific Binding (ASB) events Shi *et al.* [43] identified in the human cell line GM12878 using 14 TF ChiP-seq data sets. For heterozygous binding sites of a TF, an ASB event is defined if the number of mapped ChIP-seq reads for one allele is significantly higher than for the other allele. The authors noticed that only 19.3% of the ASB events overlap with a TFBS of the TF the ChIP-seq experiments was designed for. Hence, we only want to consider the SNPs overlapping with a TFBS of the used TF motif. Therefore, we gathered the 14 TF motifs from the JASPAR database (version 2022) [11] and computed per TF the TFBS using Fimo (version meme-5.2.0, default *p*-value cutoff) [19]. Since redundant motifs would lead to false positives later in our analysis, we clustered a combined TF motif set of the JASPAR, Hocomoco[28] and Kellis ENCODE motif [26] database using the similarity measurement and clustering approach from Pape *et al.* [37]. We checked which of the 14 TFs belong to the same cluster and kept per TF motif cluster only the TF with the highest number of associated SNPs. Doing so, we removed two TFs resulting in 368 SNPs for 12 TFs (see Supp. Sec. S3 and S5).

### 2.6 Collecting SNP-Selex data

The SNP-SELEX data was downloaded from the web portal GVAT database [49]. We gathered the SNPs called *original batch.* In total, Jian *et al.* studied 1.612.172 TF-SNP pairs for 271 different TFs. For each TF we collected all SNPs that have biological evidence to cause a differential binding event. Therefore, we filtered the SNPs for those which have an oligonucleotide binding score *p*-value < 0.05 and a preferential binding score p-value < 0.01, as proposed by the authors, resulting in 9.840 SNPs for 129 TFs. We gathered the TF motifs from Boytsov *et al.* [9], which provide optimized PWM motifs for the SNP-SELEX data set. As for the ASB data set we excluded for each TF the SNPs without a TFBS for at least one of the two alleles. If less than 5% of the SNPs associated to a TF do have a TFBS, we excluded all TF-SNPs pairs for the analysis. Next we checked which of the TFs belong to the same TF motif cluster based on the clustering used for the ASB SNPs. For each TF motif cluster we selected the TF with most SNPs, resulting in total in 33 TFs and 1.494 SNPs (see Supp. Sec. S3 and S5).

### 2.7 Method performance evaluation

We applied our method and *atSNP* to the SNP-TF pairs collected for the ASB events (see Methods 2.5) and SNP-SELEX data (see Methods 2.6). In Supp. Sec. S3 the used commands are listed. To evaluate the performance of each method, we computed a precision-recall curve and the area under the precision recall curve (AUCPR) (see Fig. 3A and B) using the R package *PRROC* [20]. To test if the area under the receiver operating characteristic (ROC) curve is significantly different between two approaches we applied the method from DeLong *et al.* [15] using the R package pROC [40].

We compared the run times of our approach and *atSNP* for 100, 500, 1.000, 10.000, 20.000 and 40.000 SNPs randomly sampled from the dbSNP database (build id 154) [42]. As both methods provide a parallel mode, we compared the runtime for 1, 8 and 16 threads (see Fig. 3C).

### 2.8 eQTL analysis

We collected the eQTLs for *EBV-transformed lymphocytes* and *cultured fibroblasts* from the GTEx portal [14] (version 8; dbGaP Accession phs000424.v8.p2) on December 21, 2022. The fine-mapped eQTLs were extracted from the file *GTEx_v8_finemapping_CaVEMaN.txt.gz* [10] resulting in 14.722 eQTLs for lymphocytes and 45.917 eQTLs for fibroblasts. Further, we downloaded the genes transcript per million (TPM) values of lymphocytes and fibroblasts from the GTEx portal *(gene_tpm_2017-06-05xν8_cells_cultured_fibroblasts.gct.gz* and *gene_tpm_2017-06-05_v8_cells_ebv-transformed_lymphocytes.gct.gz*). We computed the *D_max_ p*-value for each eQTL separately on each data set. As motif set we used a combined TF motif set from the JASPAR (version 2022), Hocomoco and Kellis ENCODE motif database. Based on the dbSNP database (build id 154), we randomly sampled 1000 SNP sets, that contain the same number of unique SNPs as the corresponding eQTL data set. Also, for each randomly sampled SNP set we computed the *D_max_ p*-value. Next, we counted over all eQTLs how often the binding sites of a TF were significantly affected *(D_max_ p*-value ≤ 0.01. We denote this count as TFcount. To identify cell type specific TFs, it is necessary to normalize the TFcount by those observed for randomly sampled data, since every motif has a different probability to occur by chance depending on the properties of the TF motif itself. Similarly, we counted for each randomly sampled SNP set how often the binding site of a TF was significantly affected. We took the mean over all randomly sampled SNP sets, denoted by bgCount. To identify TFs which are more often affected by the eQTLs of the current cell type than expected, we computed an odds-ratio as 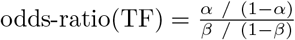 where 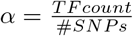, #SNPs is the number of unique SNPs per cell type and 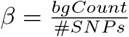.

Since we cannot distinguish TFs with binding motifs of high similarity, we want to evaluate the result on the level of TF motif clusters (see Methods 2.5). If a TF within a cluster has an odds-ratio ≥ 2 we assume it as enriched, as it is two times more often affected by the SNPs of the cell type of interest than expected by chance. For each cluster we consider as gene expression level the maximal TPM value over all TFs in the cluster. To check if the difference in expression between the TFs with enriched odds-ratio and the TFs without an enriched odds-ratio is significant, we applied an one-sided Wilcoxon rank sum test using the *wilcox.test* functionality in R.

### 2.9 Application Atherosclerosis GWAS

The atherosclerosis GWAS was download from the GWAS catalog (EFO_0003914), including the child traits *carotid atherosclerosis* (EFO_0003914) and *peripheral arterial disease* (EFO_0009783). We excluded all indels, duplicated SNPs and all SNPs from GWAS not based on an European cohort. For the remaining 255 out of 261 SNPs, we determined the SNPs in linkage disequilibrium (LD) applying SNiPA [3] using as population the European cohort and an LD threshold of 0.8. We gathered 4326 unique proxy SNPs. We computed the *D_max_ p*-value for each collected SNP associated to atherosclerosis. As input motifs we used 817 non-redundant human motifs from the JASPAR (version 2022), HOCOMOCO and Kellis ENCODE motif database. If the *D_max_ p*-value is ≤ 0.01, we assumed that the binding site of a TF was significantly affected by the considered SNP. To link the regulatory SNPs to target genes, we used 2.4 million regulatory elements (REMs) associated to target genes downloaded from the EpiRegio database [5,41]. We overlap the identified rSNPs with these REMs using bedtools’ *intersect* functionality [38].

## 3 Results

### 3.1 The differential TF binding score follows approximately the *L*(0, *b*) distribution

Consider the sequences *S*^1^ and *S*^2^ each holding an allelic variant of a given SNP and a TF model *M* that characterizes the binding behaviour of a TF. We defined the differential TF binding score (D) between *S*^1^ and *S*^2^ as the log-ratio of the *p*-values of the TF binding scores (see Methods 2.1).

To investigate the distribution of the differential TF binding scores, we decided to represent the TF models with PWMs, which are widely used and easy accessible for hundreds of human TFs, and for which other methods exist for comparison. We computed *D* for 200.000 SNPs randomly sampled from the dbSNP database for PWMs of different length. In Fig. 1 the resulting distributions of the differential TF binding scores for three TFs are visualized. For comparison, the *L*(0, 1) distribution is displayed as well. Even though we argued that theoretically *D* should be *L*(0, 1) distributed, we observed that our experiments suggest otherwise. The sequences of *S*^1^ and *S*^2^ just differ by one letter at the position of the SNP. Hence, the TF binding scores for *S*^1^ and *S*^2^ do not change much, especially if the SNP does not affect the binding site. As a result, we concluded that the corresponding *p*-values may not always be independent from each other. Consequently, we observed that *D* is close to 0 more often than one would expect for independent *p*-values. However, we empirically observed that *D* can be approximated by a *L*(0, *b*) distribution with a scale parameter *b* fitted for each TF model *M* (see Fig. 1 blue curves).

**Fig. 1.**
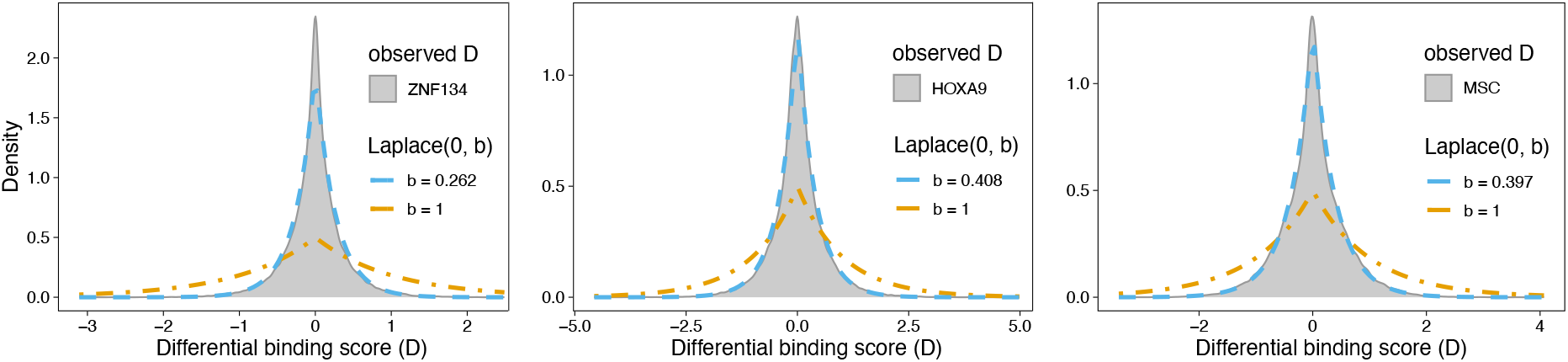
Differential TF binding score distributions for three TFs of different length. The distributions of *D* for the PWMs of TFs ZNF134 (length 22), HOXA9 (length 8) and MSC (length 10) are compared to the *L*(0,1) (orange) and *L*(0,*b*) distribution (blue). The scale parameter *b* is estimated for each TF model separately.

### 3.2 The maximal differential TF binding score distribution differs between multiple TFs

In the previous section, we assumed that we are interested in *D* for a specific sequence position, thus the sequence S and the TF model had the same length. However, that is an unrealistic assumption. Usually, the sequence is longer than the TF model. As explained in Method section 2.1, we computed *D* for all sub-sequences overlapping the SNP to identify where the binding affinity of a TF is most affected and keep the absolute maximal differential TF binding score (*D_max_*).

For 200.000 randomly sampled SNPs, we determined *D_max_* and visualized the resulting distributions for the TFs ZNF134, MSC and HOXA9 in Fig. 2A. Since the distributions are different, it is not possible to compare *D_max_* between different TFs directly. However, the usual application of such a statistic is to evaluate and compare the effect for several hundred TFs on a SNP set. To allow this comparison, we aimed to compute a *p*-value for *D_max_*. We first tried to fit known distributions to the observed *D_max_* values (using R package gamlss [39]), but this approach did not result in the same distribution for multiple TFs. As an alternative, we derived the *L_max_*(*n, b*) distribution (see Method section 2.3). The parameter *n* denotes the length of the TF model, which is given by the number of sequence windows tested by the model and the scale parameter *b* is estimated for each TF separately using the MLE from Eq. 9.

**Fig. 2.**
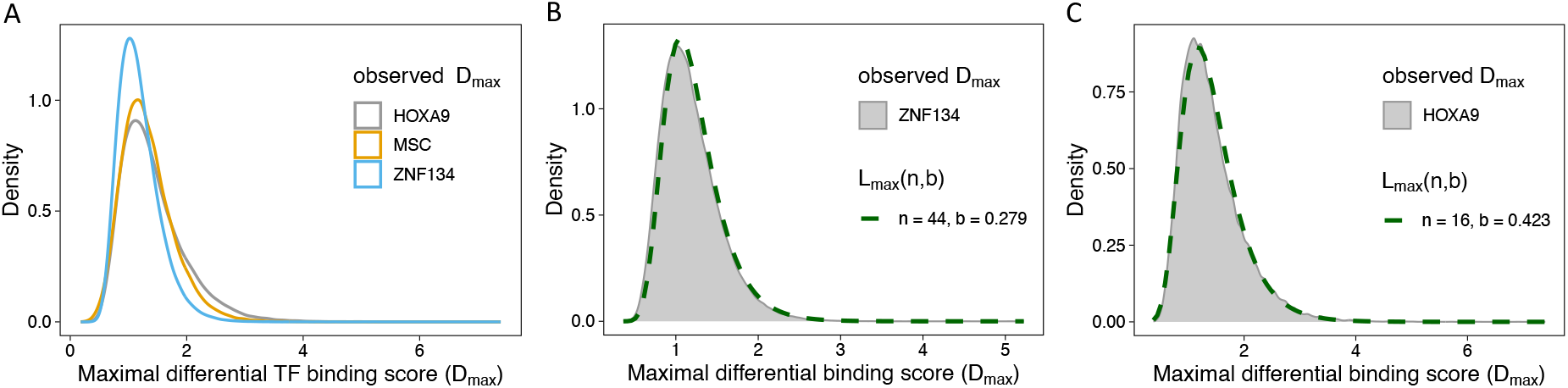
Distribution of *D_max_* values. (**A**) Density plot for observed *D_max_* values for the PWMs of TFs HOXA9, MSC and ZNF134 indicated by different colours. Distributions of observed *D_max_* values for ZNF134 (**B**) and HOXA9 (**C**) in comparison to the *L_max_*(*n,b*) distribution, parameter values for *n* and *b* (fitted) shown in the plot.

In Fig. 2 B,C we show example distributions of observed *D_max_* values for the TFs HOXA9 and ZNF134 and the corresponding *L_max_*(*n, b*) distribution. One can observe that the *L_max_*(*n,b*) distribution accurately approximates the observed *D_max_* values for each TF. Using the CDF of the *L_max_*(*n, b*) distribution we can compute a *p*-value for *D_max_* values, which are then comparable between multiple TFs.

### 3.3 Evaluation of our approach on experimentally validated TF-SNP pairs

To analyse the performance of our approach, we collected TF-SNP pairs from data sources with experimental evidence that the TF is affected by the SNP. We gathered ASB events, which are defined using TF ChIP-seq data and data collected from SNP-SELEX experiments, an *in vitro* measurement of the TF-DNA interaction strength for each allele of the SNP (see Methods 2.5 and 2.6).

To evaluate how well a method can distinguish experimentally validated TF-SNP pairs from those that are not validated, we defined a classification task, where the positive class contains the collected TF-SNP pairs. The negative class is defined as all possible combinations of considered TFs and SNPs, excluding those from the positive class. The ASB data set consists of 368 positively labeled TF-SNP pairs and 4.036 negative, and the SNP-SELEX data of 1.814 positive and 58.162 negative, respectively.

Using these data sets we do not only want to evaluate our approach, we also want to compare our method to the previously published method *atSNP* [54]. To the best of our knowledge, *atSNP* is the fastest PWM based method currently available, which also provides a *p*-value for the differential TF binding score. *is-rSNP* is too slow to be applied on the large data sets considered here [32]. We applied our approach and *atSNP* for each of the two data sets separately, and evaluated the performance for the *D_max_ p*-value in comparison to *atSNP*’s rank-based *p*-value, which is recommended by the authors. The rank-based *p*-value indicates if the log-odds ratio of the *p*-values of the TF binding score for the wildtype and the alternative allele significantly differs from what one would expect by chance. Additionally, they provided a diff-based *p*-value, that evaluates the changes in the TF binding score directly. However, the authors of *atSNP* mention that the diff-based *p*-value is not reliable, and our experiments confirmed a poor performance (data not shown).

Fig. 3 A and B show the resulting precision recall curves and the AUCPR for the ASB events (Fig. 3A) and the SNP-SELEX data (Fig. 3B). Even though the negatively labeled TF-SNP sets are several times larger than the positive sets, a reasonable AUCPR is reached for both data sets indicating the quality of our method and the rank-based *p*-value of *atSNP.* The AUCPR of the *D_max_ p*-value is improved in comparison to the rank-based *p*-value of *atSNP.* For the ASB events the AUCPR of the *D_max_ p*-value is 0.9% higher than the rank-based *p*-value. For the SNP-SELEX data set the improvement of the *D_max_ p*-value is 2.7% in comparison to the rank-based *p*-value. Additionally, for both data sets the difference of the area under the ROC curves between the *D_max_ p*-value compared to the rank-based *p*-value are significant (p-value ≤ 0.05, ASB data: *p*-value = 0.00541, SNP-SELEX data: *p*-value 9.457e^-5^) according to the method of DeLong *et al.* (see Methods 2.7).

**Fig. 3.**
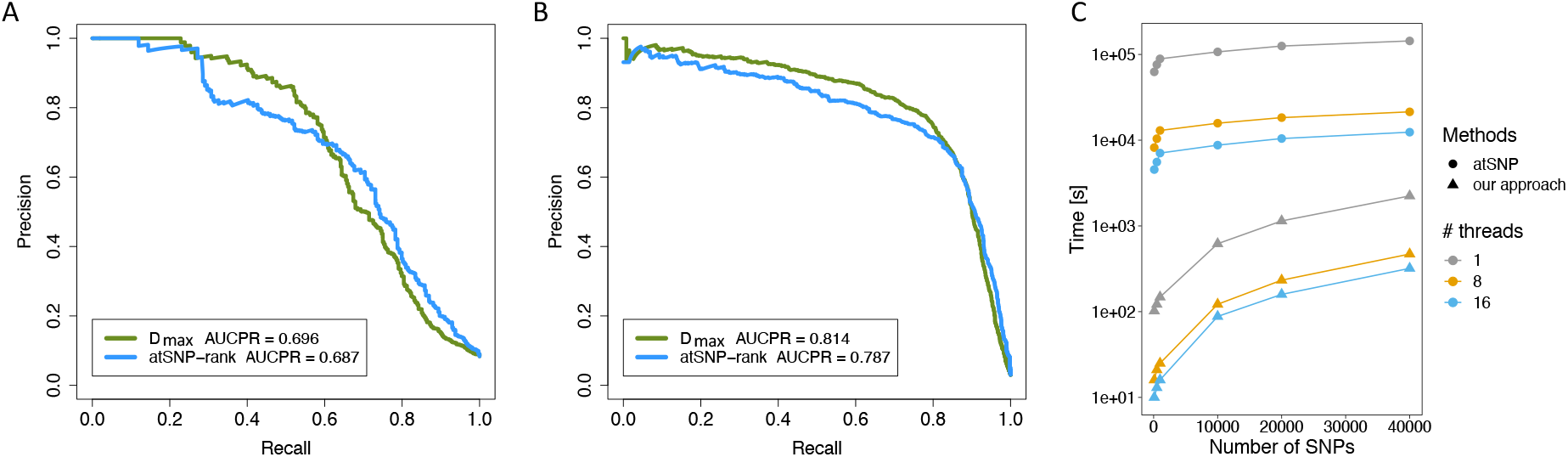
Comparison between our approach and *atSNP*. Precision-Recall curve for the ASB events (**A**) and the SNP-SELEX data set (**B**) for the *D_max_ p*-value and the rank-based *p*-value of *atSNP.* (**C**) Runtime analysis of our approach and *atSNP.* The lineplot shows the runtime for both methods (y-axis, log10-scale) for randomly sampled SNP sets of different size (x-axis) on a different number of threads.

We compared the runtime of our approach and *atSNP* for 6 randomly sampled SNP sets of sizes between 100 and 40.000 SNPs for 817 TF motifs using 1, 8 and 16 threads. As shown in Fig 3C our method is between 623 (500 SNPs on 1 threads) and 38 (40.000 SNPs on 16 threads) times faster than *atSNP.* To decide for a reasonable *D_max_ p*-value cutoff, we computed the *F*1 score for the ASB and SNP-SELEX data. The resulting *D_max_ p*-value cutoff of 0.01 is used in the following analyses.

### 3.4 Identification of TFs that are altered by genetic variants mediating gene expression

Identifying cell type specific regulators with modified binding behaviour induced by genetic variants associated to genes might be helpful to unravel regulatory pathways or molecular mechanisms. Therefore, we aimed to identify TFs more often affected by a set of SNPs than one would expect on random SNP data. Given the speed of our approach, such analyses can be done in a reasonable amount of time even for large SNP sets.

Exemplary, we analysed 14.722 eQTLs associated to lymphocytes and 45.917 eQTLs associated to fibroblasts. For each eQTL, we computed the *D_max_ p*-value for 817 human TFs. We counted per TF how often its’ binding site was significantly *(D_max_ p*-value ≤ 0.01) affected over all eQTLs. To identify TFs more often affected by the eQTLs than expected, we computed an odds-ratio between the TF counts of the eQTLs and TF counts on 1.000 SNP sets of the same size as the eQTL data (see Methods 2.8).

If the odds-ratio of a TF is > 2 we assumed that the TF is enriched since it occurs 2 times more often than expected. For fibroblasts 58 TFs and for lymphocytes 67 TFs were identified (full list per cell type in Supp. Sec. S5). Fig. 4A visualizes the odds-ratios of the analysed TFs of fibroblasts against lymphocytes, the enriched TFs with an absolute difference of the odds-ratio > 1 are labeled. For several of the cell type specific TFs, we found evidence in the literature: For instance, it has been shown that in skin fibroblasts EGR3 can upregulate genes associated with tissue remodeling and wound healing [17]. The expression of TF SNAI1 in cancer associated fibroblasts is directly associated to chemoresistance via the mediation of the extracellular matrix [18]. For lymphocytes, we identified several highly expressed TFs from the Ets-related TF family, among others, with additional evidence from the literature. For instance, in mice it has been shown that the TFs ELK1 and ELK4 function redundantly to restrict the generation of innate-like CD8+ T-cells [36]. Recently, Tsiomita *et al.* showed that ERF is a potential regulator during T-lymphocyte maturation [46]. Further, for innate lymphoid cells (ILC), which are a population of lymphocytes, it can be shown that ELK3 is regulated by the circRNA circTmem241 and that the knockdown of ELK3 significantly decreased the number of ILCs [31].

**Fig. 4.**
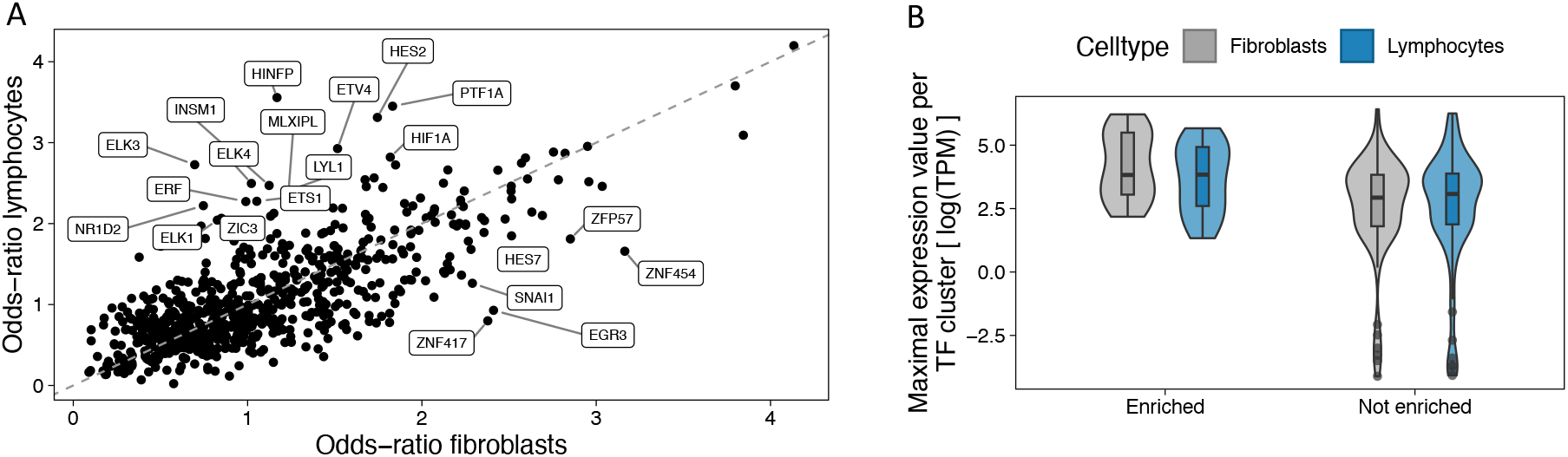
Identification of TFs that are altered by genetic variants mediating gene expression in lymphocytes and fibroblasts. (**A**) Scatter plot showing the computed odds-ratio of a TF for eQTLs from fibroblast cells (x-axis) against lymphocyte eQTLs (y-axis). The TFs are labeled if the odds-ratio > 2 and the absolute difference of the odds-ratio between the two cell types is > 1. (**B**) Violin-boxplot illustrating the differences in terms of expression values between the TFs which are more often affected by the eQTLs (enriched odds-ratio) than expected (not enriched odds-ratio) for two different cell types (coloring).

We cannot provide literature evidence for all cell type specific TFs, therefore we want to evaluate the expression values of enriched TFs in comparison to TF not enriched (Fig 4B). Since motifs from the same TF family are often highly similar, resulting in redundancy in our motif collection, we show the expression value of the highest expressed TF per cluster. According to an one-sided Wilcoxon rank sum test the differences in gene expression between the two groups are significant (fibroblasts: *p* = 0.0016, lymphocytes *p* = 0.028). Thus, many of the TFs that show strong enrichment in altered binding behaviour for eQTLs of the two cell types, are likely to mediate gene expression differences and are interesting candidates for further cell type specific investigations.

### 3.5 Identification of candidate target genes affected by the rSNPs in an atherosclerosis GWAS

Even if no eQTL data is available, one might be interested in identifying target genes that are regulated by the TFs with modified TFBSs caused by rSNPs. By combining our approach with publicly available regulatory elements (REMs), we are able to highlight candidate target genes. To illustrate this application, we have applied our approach to a set of 4326 lead and proxy SNPs of a GWAS for the disease atherosclerosis. To associate the resulting rSNPs to target genes, we used 2.4 million REMs linked to target genes downloaded from the EpiRegio webserver [5,41] (see Methods 2.9, Supp. Sec. S5).

Among the genes affected by at least one rSNP are the ABO and CELSR2 loci, which were associated with coronary artery disease (CAD) by a GWAS before [16]. CAD can be induced by atherosclerosis occurring in the large vessels supplying oxygen to the myocardium. Both loci were further implicated to play a role in lipid metabolism, with ABO, for example, being associated with total cholesterol [13] and CELSR2 representing a candidate gene at the chromosome 1p13 CAD/cholesterol locus [48]. Phenome-wide association results are depicted in Suppl. Sec. S4. The association with both, CAD and cholesterol levels, renders an involvement of hypercholesterolemia as the likely responsible intermediate phenotype. Furthermore, the binding activity of the TF OSR1 is predicted to be affected by the rSNP rs629301, which is linked to the gene CELSR2. The OSR1 gene itself was associated with the traditional CAD risk factor blood pressure [23]. The predicted functional connection of OSR1 and CELSR2 therefore might indicate an interaction between two traditional risk factors via genomic variation.

## 4 Discussion

Throughout the manuscript, we presented a new statistical approach to identify regulatory SNPs modifying the binding sites of TFs. We aimed to provide a method that allows to compute statistical significance for general TF models. We compared our new approach to the previous method *atSNP* in terms of performance and runtime. We demonstrate that our new approach is as least as accurate as *atSNP*. By comparing the runtimes of both methods, we show that our approach is extremely fast also for large sets of SNPs and hundreds of TFs.

To test our approach, we used PWMs as TF model, since they are still commonly used and available for hundreds of human TFs. Nevertheless, our approach can be applied to any other TF model once a *p*-value for the TF binding score is computed, which can be done using Monte Carlo sampling if not otherwise existing. Thus, an interesting research direction is to explore the *L_max_*(*n, b*) distribution fit to observed *D_max_* values of other approaches such as TFFMs or SLIM models.

Our approach does not directly take into account cell type or tissue specific information. However, a useful approach is to exclude motifs from TFs not expressed in the cell type or tissue of interest to reduce the number of false positive predicted rSNPs. Further, one can easily combine the predicted rSNPs with other epigenomic data as shown in the application for the atherosclerosis GWAS.

Additionally, we want to emphasize that we combined several TFs within one data set to compare the methods, resulting in a highly imbalanced data set. In our opinion, this is a more realistic evaluation setting than evaluating the TFs one by one as is often done.

Another advantage of our approach is that it has no significant additional runtime compared to widespread score-based approaches that do not assess significance. One only needs to pre-compute the scale parameter *b* for the used TF motifs. On our github repository (https://github.com/SchulzLab/SNEEP) we provide our approach implemented in C++ as an easy-to-install bioconda package, also including the pre-computed scales for the 817 TF motifs used for presented analyses.

We believe that our approach will be helpful to identify novel rSNPs and thereby contribute to the understanding of molecular mechanisms leading to various traits and diseases.

## Acknowledgment

We thank the GTEx Portal and the EBI GWAS catalog for providing the data used in the applications and Fatemeh Behjati Ardakani for proof-reading.

