## SupplementaryData for "A statistical approach to identify regulatory DNA variations"

January 31, 2023

#### Supplementary Section S1. Fit the scale parameter $b$

In all analyses we conduct within this work we used as TF motif set either the collection of non-redundant human PWMs combined from JASPAR (version 2022), Hocomoco and Kellis ENCODE motif database or subsets of it. We removed flanking bases of the TF motifs with an entropy higher than 1.9, since we observed that TF motifs with flanking bases that exhibit a high entropy have a negative effect on the fit of the distribution to the observed $D_{max}$. To apply our method, we pre-computed the scale parameter $b$ for each TF motif. To do so, we randomly sampled 200.000 SNPs from the dbSNP database (build id 154). For each TF, we computed $D_{max}$ for all SNPs. These values are plugged in the MLE of the PDF of $L_{max}$ and numerically solved using Newton's method to approximate $b$ (done with python library scipy.optimize Virtanen et al. [2020]). We want to describe the tail of the distribution of $D_{max}$ as accurate as possible. Therefore, we minimized the mean squared error (MSE) for the tail of the distribution of $D_{max}$ (25% of all values) by decreasing / increasing the estimated scale parameter $b$ by 0.01 as long as the MSE decreased.

#### Supplementary Section S2. Details how to compute the exact TF binding score distribution

We used PWMs as TF model. Therefore, we downloaded the TF models in TRANSFAC format from the JASPAR, HOCOMOCO and Kellis ENCODE motif database. We convert the motifs to Position Weight Matrices (PWMs), thereby we added an epsilon of 0.001 to every entry to avoid 0 entries. To apply the dynamic programming approach of Beckstette *et al.* [Beckstette et al., 2006], we shifted the resulting log-likelihoods in such a way that all values are $> 0$ and rounded them with an accuracy of 0.001. We precomputed the exact TF binding score distribution for all given PWMs (for more details see *Section Calculation of exact PSSM score distributions* in the paper of Beckstette *et al.*). An implementation of their approach can be found on our github repository (https://github.com/SchulzLab/SNEEP), see file *src/pvalue_copy.hpp.*

### Supplementary Section S3. Used commands to apply our approach and *atSNP*

For a better reproducibility of our results, we provide all executed commands and the input files (see Supp. Sec. S5). For all analyses we used as genome version hg38, beside for the SNP-SELEX data where we used hg19. The dataset specific input files are indicated with <>. The SNP and motifs data for the different data set for our approach and *atSNP* are provided in their required format, the content is the same.

#### Used commands to apply our approach

In the following the used commands to run our approach are listed. All script and more details how to run them can be found in our github repository: https://github.com/SchulzLab/SNEEP.

As a first step, we need to estimate the scale parameter *b* for the used PWMs once using

```
bash estimateScalePerMotif.sh 200000 <motifs> <outputDir>
<motifNames> 1.9
```

As *motifs* we used the file *combined_Jaspar_Hocomoco_Kellis_human_transfac_jaspar_2022.txt*, which can be obtained from our ZENODO data repository. *motifNames* lists the names of all TF motif for which we wish to compute *b*. The result is a file providing for each considered TF model the estimated scale parameter *b*, called *estimatedScalesPerMotif_1.9.txt* in the following.

To compute the results for the ASB events, the SNP-SELEX data and the runtime analyses we used the following command:

```
time ./src/differentialBindingAffinity_multipleSNPs -o <outputDir>
-n <numberCores> -p 1.0 -c 1.0   -s estimatedScalesPerMotif_1.9.txt
-j 0 <motifFile> <input-snps>  <path-to-genome-file >.
```

The file *motifFile* can be found in our ZENODO data repository as well as the *input-snps*:

- ASB events: snps_ASB.txt, motifs_ASB.txt
- SNP-SELEX data: snps_SNP_SELEX.txt, motifs_SNP_SELEX.txt
- the randomly sampled SNPs can be found in the files sampledSNPs100.txt, sampledSNPs500.txt, sampledSNPs1000.txt, sampledSNPs10000.txt, sampledSNPs20000.txt, sampledSNPs40000.txt, motifs: *combined_Jaspar_Hocomoco_Kellis_human_transfac_jaspar_2022.txt*

For the eQTL analyses we performed a background sampling which can be done automatically using:

```
time ./src/differentialBindingAffinity_multipleSNPs -o <outputDir>
-n 16 -p 0.5 -c 0.01 -s estimatedScalesPerMotif_1.9.txt -j 1000
-l 10 -k dbSNPs_sorted.txt
combined_Jaspar_Hocomoco_Kellis_human_transfac_jaspar2022.txt
<input-snps>  <path-to-genome-file >
```

The *input-snps* we downloaded from the GTEx Portal (see Methods 2.8), the file *dbSNPs_sorted.txt* is part of our github repository and the file *combined_Jaspar_Hocomoco_Kellis_human_transfac_jaspar_2022.txt* can be found in our ZENODO data repository

To link rSNPs to regulatory elements as we did it for the atherosclerosis GWAS, we executed the command:

```
time ./src/differentialBindingAffinity_multipleSNPs -o <outputDir>
-n 10 -p 0.5 -c 0.01  -r REM.txt -s  estimatedScalesPerMotif_1.9.txt
-g ensemblID_GeneName.txt  -j 0
combined_Jaspar_Hocomoco_Kellis_human_transfac_jaspar2022.txt
<input-snps.txt> <path-to-genome-file>
```

The files *interactionsREMs.txt* and *ensemblID_GeneName.txt* can be found in our github repository and in our ZENODO data repository the SNPs of the GWAS atheroclersis are stored in the file snps_atherosclerosis.txt and the motifs are provided in the file *combined_Jaspar_Hocomoco_Kellis_human_transfac_jaspar_2022.txt*.

##### Commands used to run *atSNP*

To run *atSNP* (version 1.14.0) the following commands where executed in R:

```
pwms <- LoadMotifLibrary(<motifFile>, tag = "MOTIF", skiprows = 2,
skipcols = 0, transpose = FALSE, field = 2, sep = " ", pseudocount = 0)

snps <- LoadSNPData(filename = <snpFile>, genome.lib = <genomeVersion>)

scores <- ComputeMotifScore(pwms, snps, ncores = <numberCores>)

diffBind <- ComputePValues(motif.lib = pwms, snp.info = snps,
motif.scores = scores$motif.scores, ncores = <numberCores>)
```

The used motif input files and the SNP files are uploaded to our ZENODO data repository:

- ASB events: snps_ASB_atSNP.txt, motifs_ASB_atSNP.txt
- SNP-SELEX data: snps_SNP_SELEX_atSNP.txt, motifs_SNP_SELEX_atSNP.txt
- randomly sampled data: snps: sampledSNPs100_atSNP.txt, sampledSNPs500_atSNP.txt, sampledSNPs1000_atSNP.txt, sampledSNPs10000_atSNP.txt, sampledSNPs20000_atSNP.txt, sampledSNPs40000_atSNP.txt, motifs: motifs_JASPAR_HOCOMOCO_Kellis_1.9.meme

As *genomeVersion* we used "BSgenome.Hsapiens.UCSC.hg38" for the ASB events and "BSgenome.Hsapiens.UCSC.hg19" used for the SNP-SELEX data.

#### Supplementary Section S4. Identification of candidate target genes affected by the rSNPs in an atherosclerosis GWAS

Phenome-wide association results are depicted in Fig. 1 at the end of the document.

#### Supplementary Section S5.

We provide a ZENODO data repository (https://doi.org/10.5281/zenodo.7588272) with the following content:

- ASB events and the corresponding PWMs used to evaluate our approach (for our approach and *atSNP* in their required format, for details see Supp. Sec. S3).
- result of our approach and *atSNP* for the ASB events (data for Fig. 3A) (ASB_result_sneep_atSNP.txt)
- SNP-SELEX data used to evaluate our approach (for our approach and *atSNP* in the required format, for details see Supp. Sec. S3).
- result of our approach and *atSNP* for the SNP-SELEX (data for Fig. 3B) (SNP_SELEX_result_sneep_atSNP.txt)
- files containing the enriched TFs (odds-ratio > 2) for lymphocytes (enriched_TF_lymphocytes.txt ) and fibroblasts (enriched_TF_fiborblasts.txt )
- input SNPs of the GWAS atherosclerosis (snps_atherosclerosis.txt) and the result of our approach (atherosclerosis_result.txt)
- TF input motifs in TRANSFAC format for our method and in MEME format for atSNP (see Supp. Sec. S3)

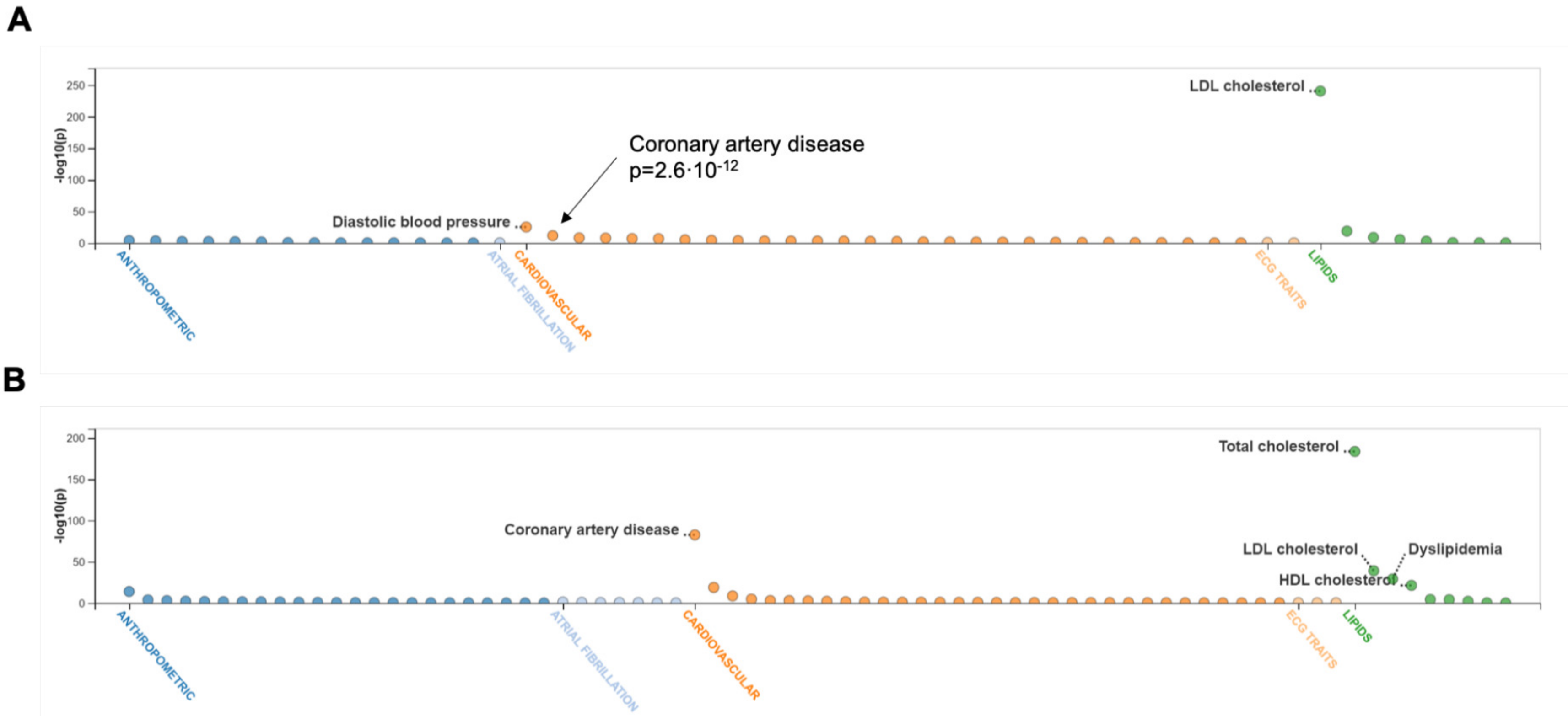


**Figure 1:** Phenome-wide associations of SNPs at predicted TF binding sites. **A** Rs582094 at the ABO CAD locus. **B** Rs629301 at the CELSR2-SORT1 CAD locus. Only significant interactions are shown ($p \leq 0.05$). Data extracted from Cardiovascular Disease Knowledge Portal (Accessed on 2021-09-01; `https://cvd.hugeamp.org/variant.html?variant=rs582094`; `https://cvd.hugeamp.org/variant.html?variant=rs629301`).
